# Discovering footprints of evolutionary chromatin response to transposons activity: merging biophysics with bioinformatics

**DOI:** 10.1101/513572

**Authors:** S. Vitali, E. Giampieri, S. Criscione, C. Sala, I. Faria do Valle, N. Neretti, G. Castellani

**Author notes:** these authors contributed equally to this work (S. Vitali, E. Giampieri).

## Abstract

Transposons are genome components that account for the majority of genome size in many organisms, behaving as parasitic entities and interfering with the translation mechanism. Chromatin structure influences the activity of transposons, by coordinating genome accessibility for the expression and insertion of these sequences. As a case study, we show evidences of an evolutionary response of the chromatin structure to a variation in the activity of Long Interspersed Elements (LINEs) during mammals evolution, with focus on the murine radiation and primate evolution. LINEs activity was measured using a biophysical approach for modeling LINEs as an ecosystem, where different strains of transposons might reproduce, die and compete for access to the translational machinery of the host. The model, based on the discrete stochastic processes of amplification and deactivation of LINEs copies, has been adapted to the data using Bayesian statistics to estimate its main parameters: rate of growth of transposons copy number and rate of past competition between transposons variants. This approach allows to estimate the activity of ancient LINE strains still present in the genome as deactivated components, and the possible competition among different strains. We leverage these results to highlight how the change in the chromatin structure of the murine species seems to be following an increase of LINEs activity during the appearance of the murine specific strain Lx. On the contrary, a similar response is absent in primates evolution, which follows a decrease of LINEs activity during the amplication of primate specific LIMA/LPB strains.

## Introduction

Transposable elements (TEs) are protein coding DNA sequences that can move from one genomic location to another and increase their copy number in the host-genome as parasitic entities. They constitute a large portion of many species’ genomes (e.g., 45% of the human genome), and a large variety of TEs sequences is annotated, subdivided into two classes on the basis of the transposition mechanism: DNA-transposons - involving a DNA copy and paste mechanism, and retro-transposons - involving RNA-mediated transcription mechanisms [9]. By following a taxonomic approach, each class is further partitioned into sub-classes, families, subfamilies and elements, on the basis of their structural organization, the proteins they encode, and the sharing of specific insertions, deletions or substitutions [42].

The interaction between the host organism and TEs has been often described as a host-parasite relationship because TEs cannot perform their transposition activity out of the host organism and such activity has in general a negative impact over the host fitness, due to deleterious insertions in the genome of new TEs copies. The impact over the host fitness of TEs appearance and amplification has been extensively investigated to study how TEs copies reach fixation in host populations and how TEs abundances are distributed [7, 41, 12, 1, 33, 34, 32]. However, such host-parasite interaction is peculiar because the survival of the TEs is often subordinated to the survival of the host at least in terms of population, especially for TEs sub-types which are incapable to transfer from one host to another (*horizontal transfer*). Thus, the loss of fitness due to TEs activity is also responsible for the appearance of host-driven regulation (epigenetic silencing [**Liu**]) and self-regulation (evolution of traits constituting a selective disadvantage at the copy level as for example a lower transposition rate [20, 32]), to limit the number of new TEs insertions and their impact over the genome functionality [**Sun 2018**, 5, 22].

Phenomena related to the molecular nature of TEs may also occur, for example mutations, insertions and sequence rearrangements, which may lead to functional variations of the elements. Molecular events can lead to the formation of new kinds of TE or the inactivation of all the copies of a specific variant, thus preventing any new duplication.

Beside sequence mutation, the inactivation of all the remaining TE copies with intact DNA sequence is induced by epigenetic variations of the host genome that silence transposition activity [**Liu**, 31], including DNA methylation, chromatin remodeling, and miRNAs.

The configuration of the chromatin is fundamental to regulate the accessibility of genome portions to transcription enzymes. Such transcription enzymes are necessary to initiate a transposition event, and their ability to reach the site of TEs insertions has an impact over TEs copy number and transposition dynamics, especially for the retroelement class [13]. Therefore, we distinguish two wide types of chromatin configuration for each copy: *closed chromatin configuration* or *heterochromatin*, not accessible by transcription enzymes, and *open chromatin configuration* or *euchromatin*, accessible by transcription enzymes.

The interdependence between TEs community and the host genome, together with the replication mechanisms of the elements and the possibility of appearance and disappearance of TEs varieties, suggests a strong parallelism between TEs dynamics in the genome and population dynamics in a ecosystem [42, 37]. Both the niche theory and the neutral theory of biodiversity [4] have features convenient to describe TEs ecosystem. Niche theory is based on the partitioning of resources between competing species that occupy different ecological niches [8], while in the neutral theory the main hypothesis is the equivalence of species and that stochastic mechanisms (such as demographic stochasticity, migration, and speciation) are the main forces shaping the community [21, 4]. Therefore, neutral theory is mainly applied to describe species at the same trophic level or that occupy the same niche.

However, TEs ecosystem contains some peculiarities [42] that needs to be explicitly addressed in the modeling: 1) TEs create and continuously reshape their own environment, because TEs copies constitute a large part of the genome landscape in which new copies may insert without deleterious effect on the cell functionalities; 2) the natural selection acts on two levels, the genome level and the host one. In this regard, we distinguish *transposon ecology*, describing the interaction of TEs with the genome and cellular environment only, from *genome ecology*, which includes the interaction with the external environment mediated by the host fitness [28].

From a transposon ecological perspective a single copy could be considered as an individual, and all the individuals belonging to the same family or subfamily as constituting a species, meaning that they occupy the same ecological niche in the genome and can be considered equivalent species [42]. In the present work we reserve the word *species* to the element level for modeling purposes. Therefore, different elements belonging to the same family or sub-family are treated as different species, each one with its own number of copies. The copy number of an element constitutes the abundance, or the number of individuals of that species. The community is composed by all the TE species belonging to the same family or subfamily, which can be located at the same trophic level.

Here we focus on the study of a particular family of *non-long terminal repeats*: the Long Interspersed Elements (LINEs), which are the most abundant family of TEs in mammals, in terms of biomass and belongs to the retro-element class because of their transposition mechanism (see Box 1).The LINE community in mammalian genomes is mainly composed by defective and/or silenced copies of inactive elements that reached fixation in the genome. LINEs often evolved on a single lineage, in particular in primates [26], with a subsequent appearance of active elements, making competition between different elements negligible. Coexistence of multiple LINEs lineages is documented for ancient LINEs [38] and currently in mouse [29], where LINEs frequently recruited novel 5’UTR sequences [40], suggesting that simultaneous activity of non-homologous promoters does not introduce a competition between the elements. The stratification of such elements in the genome, due to the absence of horizontal transfer, will be used in this work to infer the changes in time that may have occurred in the dynamics of LINEs.

### Box 1. LINEs transposition mechanisms

LINEs family belongs to the retro-elements class, which means that their replication is RNA mediated (Figure 1B. LINEs are incapable of horizontal transfer [1] therefore fixation of new copies takes place mostly along the germ line. The replication mechanism has low fidelity and two types of copies can be distinguished: full-lengths, capable of autonomous transposition, and defectives, which depend on full-length copies molecular machinery for transposition.

**Figure 1:**
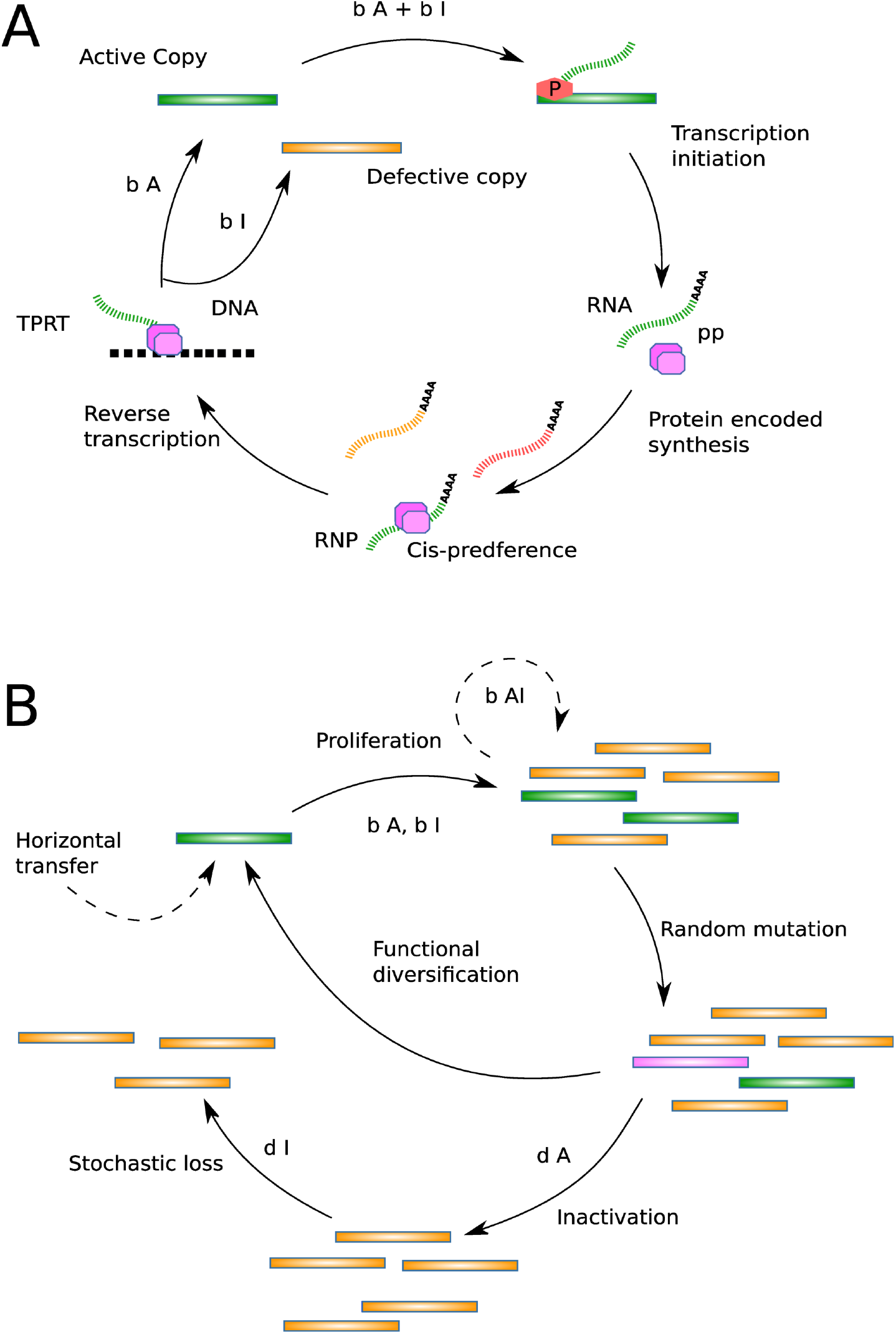
LINEs transposition and activity cycle diagram. A Diagram of the birth-death process of defective (orange) and full-length copies (green) from a full-length master copy by retro-transposition and B cycle of LINE amplification in the genome. Random sequence mutation may generate a new element species (pink) which start the cycle again and eventually become a competitor. Dashed lines describe negligible processes.

Full-length elements contain a promoter region (5’UTR), two protein coding regions (ORF1, ORF2) and a poly-A tail (3’UTR). The internal promoter directs transcription initiation, and permits autonomous transposition. When the transcribed RNA reaches the cytoplasm, the protein encoding regions ORF1 and ORF2 are translated to an RNA-binding protein and a protein with endonuclease and reverse-transcriptase activities, respectively. Both proteins show a strong cis-preference; consequently, they preferentially associate with the RNA transcript that encoded them to produce what is called a ribonucleoprotein (RNP) particle. After coming back into the nucleus, the proteins on RNA can open a nick in DNA and produce a DNA copy of the template through a process termed target-primed reverse transcription (TPRT). The new insertions often result in low fidelity copies of the parent LINE [10]. Some transposition events are incomplete such that the inserted copy is incapable of autonomous retrotransposition; for example, L1 insertions are often 5’-truncated (e.g. Figure 6B of [11]). Furthermore, a transcribed incomplete copy can hijack the retro-transposition machinery of autonomous copies to duplicate into a new location: a process called trans-complementation. The phenomenon of trans-complementation has been observed, for example, in LINE-1 retro-elements, although it should happen at a much smaller rate than retro-transposition in *cis* [**Wei2001**].

The genome environment (chromatin state, genome location, etc.) is unique to each of the TEs copies. Thus, full-length LINE copies may differ in their level of transposition activity [6, 36], and the stochasticity at the individual level could have a significant impact on the structure of the entire community, supporting the neutral approach to describe the community dynamics.

These peculiarities of LINEs replication mechanism and their evolutionary history inside mammalian genomes support our assumption that LINEs community in mammals could be successfully described by a birth-death process (see Box 2) under competitive neutrality hypothesis. Competitive neutrality represents the absence of direct competition among different LINE species, which can be considered all equivalent. Thus, all the copies of all elements in the community could be characterized by the same transposition activity, sequence divergence, and death rate [42].

### Box 2. Birth-Death processes and Master Equation

Birth-death processes are used to describe changes in the abundance of a population, whose dynamic can be represented by a discrete stochastic increase (birth) and decrease (death) of one individual at the time. They are used to describe real populations, such as animals or plants in an ecosystem, or chemical reactions inside a cell, such as the numbers of copies of a specific protein.

For discrete stochastic processes such as these, the natural description is the Master Equation (ME) [25]. ME describes the probability of observing the system in one of the possible states representing the number of individuals given the knowledge of the state of the system in the past. Birth and death processes correspond to the transitions between different states.

Under reasonable assumptions about the birth and death probability rates the system will reach a stationary state in time, which means that the probability of observing the system in any individual state (a certain number of individuals) will quickly converge to a fixed value, independently from the starting state. This means that the master equation can be solved in a general way, obtaining what is called the stationary distribution.

If we describe with *P_n_*(*t*) the probability to observe the system in the state *n*, i.e. to observe *n* individuals, at time *t* and we characterize the birth death process with *B_n_*, the time independent transition probability from state *n* to *n* + 1 due to the birth of a new individual, and *D_n_*, the time independent transition probability from state *n* to *n* − 1 due to the death of an individual, the resulting equation for the evolution of the population probability distribution can be written as:

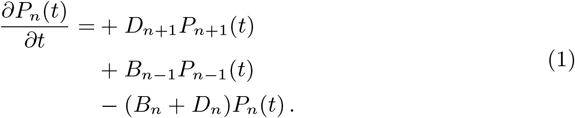

Therefore, the probability of observing a certain number *n* of individual can increase due to the chance of the system being in the state *n* + 1 and observing one individual death, or being in the state *n* − 1 and observing one individual birth. It can also decrease over time if individuals are born or died (moving the system respectively to the state *n* + 1 and *n* − 1). When these contributes are balanced such probability does not change over time and the stationary state is reached: 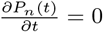.

As previously mentioned, we restrict the model to LINE family only, for discarding possible interaction with different TEs entities. Moreover, we reduce the variability of the *abiotic* component (in this context genes, repetitive sequences, and intracellular components) of the ecosystem by including only mammalian genomes in the study.

We model LINEs copy number distribution inside a cohort of mammalian reference genomes under the hypothesis of competitive neutrality [27] through a Master Equation approach [3, 25] (see Box 2) by which the Relative Species Abundance (RSA, see Box 3) can be described by a negative binomial distribution. The negative binomial distribution depends on two parameters: the “probability of success”, *x*, and the “number of failures”, ϒ. The value of such parameters allows to determine if equilibrium assumption - between the host and LINEs activity - holds for the LINEs ecosystem. In this work we use the value of ϒ to study whether the LINEs ecosystem could be treated as if in equilibrium: if the system is out of equilibrium we expect to observe an RSA following a negative binomial distribution with ϒ ~ 1; instead, if equilibrium holds, we expect ϒ ≫ 1 (see Supplementary Material A.1).

We consider that simultaneous activation of elements sharing the same promoter may introduce a disadvantage for the species that compete for the translational molecular machinery. According to our model a competitor could appear, with a certain probability, every time a new active copy is created in the system. The occasional competition between two LINEs species induces a reduction of the birth rates of the competing entities, breaking spontaneously the hypothesis of competitive neutrality. Thereafter, the extinction of one of the competitors restores the neutrality. The two models, competition and neutrality, result to be nested in their parameter description (see section methods for details). Here we will test the two models by fitting the RSA of LINEs communities through a hierarchical Approximate Bayesian Computation (ABC) method and determining the most appropriate model based on the approximated Bayesian factor. Moreover, we will apply a sliding window analysis to study the evolution of the RSA in time together with the chromatin state characterization of the individual copies. This analysis will allow to estimate the concomitance of changes in chromatin status and dynamics rates of the population.

### Box 3. Relative Species Abundance (RSA)

Relative Species Abundance (RSA) refers to the abundance of each species (the number of individuals belonging to that specie at the moment of sampling) in relation to the size of the ecosystem (total number of individuals). It is often applied to describe the variety of species occupying the same trophic level, such as all the tree variety in a forest.

A common representation of the RSA is the frequency histogram, where the species are clustered based on the number of individuals (usually binned using log 2 divisions and called Preston plot).

This description corresponds to the probability distribution of observing a specific number of individuals for each species in the ecosystem.

In the case of LINEs ecosystem the number of individuals corresponds to the copy number of the element and the species to the LINE species.

Under neutral and stationary assumptions the observed RSA frequency histogram corresponds to a possible realization of the stationary distribution of a single species ME correctly describing the population dynamics.

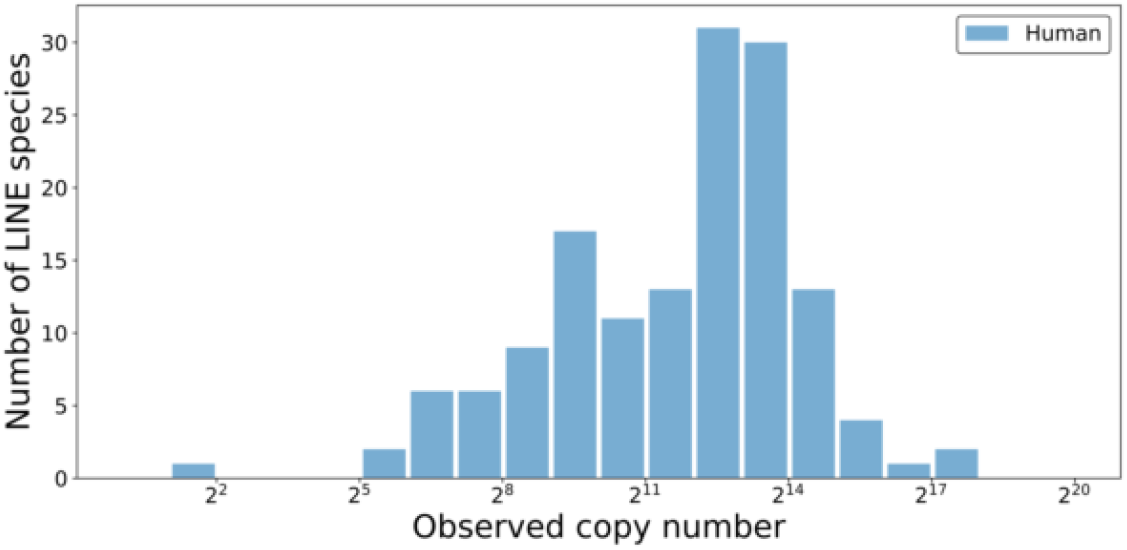
RSA of LINEs in Human genome.

## Methods

### Data sources

LINE abundances were calculated using RepeatMasker annotation (http://www.repeatmasker.org) [39] for human genome build hg19 and 45 other mammalian species (see Supplementary Tables B1, B2, B3). LINE consensus sequences were downloaded from RepBase [23, 24] (http://www.girinst.org). Chronological ordering of LINEs in human, chimp, rhesus macaque, mouse and rat genome was derived from genome wide defragmentation results [19]. Chromatin structure data are available for mouse [45] and human [15]. The employed chromatin state assignment was conducted by using ENCODE chromatin models from the ChromHMM method [14].

### Chromatine state assignment

Chromatin structure data were used to assign each LINE copy to open or closed chromatin state based on their location (coordinates) in the reference genome. Open and closed chromatin states were defined according to the available reference for mouse [45] and human [15] for the germ line. Genome regions for whom both open and closed state annotation were present and overlapping were classified tentatively as “weakly open”, and this classification was then used for the LINEs located in these regions.

We tested grouping weakly-open chromatin population with both open chromatin and closed chromatin to assess if this choice affected our results, but did not observed any significant difference. Thus, the weakly-open and the unknown state have been included in the closed chromatin group, which encloses most of the LINE copies, and thus will less likely to be affected by this approximation.

### Sliding window analysis

LINEs community in human, chimp, rhesus macaque, mouse and rat genomes have been sorted by their relative time of appearance and amplification in the genome according to specific host genome analysis [19] (different genomes may show minor variations of this ordering). By dividing the ordered group of LINEs into intervals (windows), each of them containing a fixed number of elements, a different realization of the LINEs ecosystem is obtained. By considering different windows (obtained by sliding the window along the timeline), a different sub-sample of elements active in a distinct evolution stage of the genome can be selected, representing a picture of the LINE community in a different evolutionary stage of the genome. The sequence of the windows allows to analyze the changes in time of the LINEs community in terms of the variations of: the RSA distribution patterns (see Box 3); the average percentage of insertions in open chromatin regions; and the average abundance. We used windows length of *N* = 15 number of LINEs species, sorted by their estimated order of appearance in the host genome. Windows lengths of *N* = 30, 40 produce consistent results, with smoother temporal trends.

### Stochastic model

The dynamics of the abundance inside the genome of LINEs species is described using a two-dimensional birth-death Master Equation (see Box 3). This Master Equation models the LINEs system in terms of the number of active copies (*n_A_*) and the number of inactive ones (*n_I_*), and includes the processes that change their number over time (duplication, inactivation, etc.) depicted in Figure 1a-b.

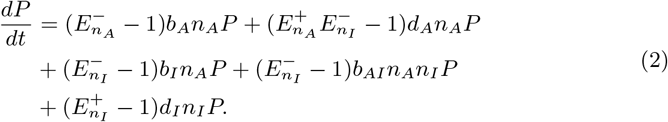

where we consider *P* ≡ *P* (*n_A_, n_I_, t*) and the Van Kampen step operators [25] 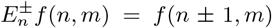, the lower index *n* indicates the variable on which the operator acts, the upper index + or determines the direction of the unitary change of the value of the variable *n*.

The variables *n_A_* and *n_I_* represent respectively the number of full-length copies (autonomous in self replication) and the number of defectives copies (non-autonomous or inactive) of a specific element in a genome. Each term in the right-hand side of equation 2 represents one of the stochastic process examined to define LINEs dynamics: *b_A_, d_A_*, rate of birth and death of full-length copies; *b_I_, d_I_*, rate of birth and death of defective copies; *b_AI_*, rate of birth of defective copies by trans-complementation.

The marginal stationary distribution *P* (*n_I_*), that will be referred to as *P_nI_*, corresponds to the RSA of the LINE species, which represents the probability to observe a species with a certain number of individuals (the copy number) inside a community (the genome), as the inactive copies are the one that are observed in the genome.

When equilibrium is reached for both active and inactive copies, the RSA corresponds to a negative binomial distribution. In fact, taking *n_A_* as a constant, the equation 2 for *n_I_* describes a well-known ecological neutral model, successfully applied to several real ecosystems [44, 35]. If equilibrium for active species does not hold (*b_A_* ≪ *d_A_*) (the LINE species tend to go extinct), we can still obtain a distribution of the values to which *n_I_* converges after the extinction (obtained when the “absorbing state” is reached, i.e. there are no more active copies and the specie is extinct and *n_A_* = 0). Thus the RSA distribution is composed by multiple replicas of the stochastic process and this distribution converges to a negative binomial distribution. The convergence to a negative binomial distribution requires the excision and transcomplementation processes to be neglected. The two regimes, with the number of active copies in or out of equilibrium, can be distinguished because the expected value of the parameters are different due to biological considerations (see Supplementary Material Section A.1).

A spontaneous breaking of the competitive neutrality assumption is introduced by assuming that contemporary activation by the same promoter region of two LINEs reduces the birth rates of both active (*b_A_*) and inactive (*b_I_*, *b_AI_*) copies in equation 2 by a factor *n*_1_*/*(*n*_1_ + *n*_2_), where *n*_1_ and *n*_2_ are the full-length copies of the two competing elements. This reduction is due to the competition for the polymerase complex between the two species triggered by shared promoter regions. A smaller birth rate will result in a lower abundance for the competing LINE species, in comparison to the elements not affected by the competition mechanism, and will induce deviations from the expected distribution by generating a bimodal behavior (one mode for each kind of process). We surmise that the distribution arising from this type of competition is a mixture of two negative binomials (equation 3) for a range of parameters compatible with real data (confirmed by numerical simulation, see Supplementary Figure A4). This distribution can therefore be expressed as the weighted combination of two negative binomial distributions:

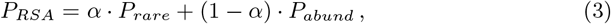

where *α* is the mixture coefficient, related to the probability that two elements compete, and with *P_rare_* and *P_abund_* representing respectively the RSA of rare and abundant elements (those with competition and those without, see Supplementary Material Section A.2).

The negative binomial distribution depends on two parameters: the “probability of success”, *x*, and the “number of failures”, ϒ:

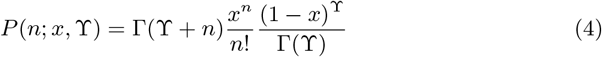

The expectation value of the copy number for the LINEs RSA is defined by the parameters of the negative binomial distribution by the relation:

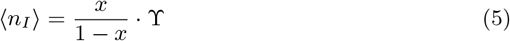

where, in the case of competition, the parameters of the distribution of the abundant species, which contains the majority of the LINE species, can be employed.

### Approximate Bayesian Computation

To estimate the parameters *x* and ϒ from the RSA data, an Approximated Bayesian Computation approach (ABC) was used. Using this approach one defines a prior distribution over the parameters, and generates a posterior distribution over their values including the information contained in the data: (i) a set of parameters is taken from the prior distributions to build a negative binomial or a mixture of two negative binomials distribution representing the RSA distribution; (ii) a sample of abundances is generated according to such distribution; (iii) the distance between the generated data and the observed data is computed (see Supplementary Material Section A.3 for details).(iv) the set of parameters taken from the priors is accepted if such distance is under a given threshold; (v) the procedure (i-iv) is repeated ~ 10^6^ times and each accepted set of parameter is stored: this collection represent an approximated sampling from the posterior of the parameters.

The final (approximated) posterior distribution of the parameters represents the probability that a certain set of parameters describes the data according to the model.

To study the variation in time of the RSA in the sliding windows analysis each window RSA was fit by such ABC method. Thereafter, each window’s RSA was represented by the average of the posterior distributions obtained for each parameter. To overcome the small sample size of window’s RSA, we employed informative priors (instead of weakly-informative ones) estimated using a hierarchical method (see subsection “Hierarchical approach for prior estimation”).

Model selection was achieved comparing the posterior probability of each model, estimated computing the ratio between the number of accepted parameters sets over the total number of simulated parameters. Specifically, for each pair of models we computed the logarithm of the ratio of the two posterior probabilities of each sample.

This method computes an approximation of the Logarithmic Bayes Factor, defined as the log ratio of the probabilities that each model is the actual true model, without incorporating prior beliefs about the plausibility of each mode [2].

This method is akin to a Bayesian equivalent of the likelihood ratio tests such as BIC (Bayesian Information Criterion) and AIC (Akaike Information Criterion) [**Marin2012**], but including an implicit penalization for the number of parameters that each model possesses and the shape of the prior for each parameter: wider priors (encoding for less certain parameters) cause a greater penalization than narrower ones (encoding for more information available about them).

### Hierarchical approach for prior estimation

The definition of the prior distribution over the parameters is a critical step in Bayesian computation. Given the relationship between the observed data (the 46 mammalian genomes), there is a similarity in the estimated properties of the distribution of RSA. This similarity would not be sufficient to include all the data in a single analysis, but it would be incorrect to ignore it as well.

This can be solved using a hierarchical approach, where the priors are modeled using hyper-priors that represents the similitude between the data and can be estimated from the data themselves. Given the high dimensionality of the data at hand, and the low acceptance rate even for simple cases, the power prior method was employed. Using this technique the analysis was performed twice for each sample by applying the same ABC method described in the previous sub-section: once with weakly-informative priors, and once with priors built from the combination of the posteriors of the first analysis of all the 46 mammalian genomes. Such combination is the mixture of all the obtained posteriors with equal weights (same number of accepted parameters), indicating the general prior for a mammalian genome.

The posterior for each mammalian genome LINEs RSA obtained in the second phase of the analysis has been employed as prior in the sliding windows RSA analysis of the same genome.

### Numerical simulations

To test if the dynamical model can generate a negative binomial distribution beyond the given assumptions, we performed numerical simulations. We used the Stochastic Simulation Algorithm (SSA) [17, 18] for the active copies dynamics, while for the inactive copies dynamics we used the tau-leap algorithm for the case *b_AI_* = 0 and a hybrid algorithm when trans-complementation is considered.

The hybrid algorithm consists in the estimation of the expected number of inactive copies by ordinary differential equation (ODE) numerical integration at the beginning of each time interval 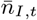. This algorithm was chosen instead of the pure SSA to reduce the computation time (SSA could take an infinite time to converge for exponential growth systems). Then, tau-leap algorithm was applied to generate a stochastic increment associated to the time interval Δ*n_I,t_*. The number of inactive copies at the end of the time interval is thus determined by the sum 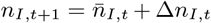. Comparison to the theoretically correct Gillespie algorithm was performed to test the accuracy of the hybrid simulation method. Simulations of the competition mechanism in both regimes (*b_AI_* = 0 or *b_AI_* > 0) were performed to check if the solution was compatible with a mixture of negative binomials, for both equilibrium (*b_A_* ≈ *d_A_*) and out of equilibrium (*b_A_* ≪ *d_A_*) conditions (see Supplementary Material Section A.4).

## Results

### The competition model is preferred over the neutral model for the majority of the RSA

The sliding windows of the RSA distribution patterns have been fitted by a negative binomial and by a mixture of negative binomials through the ABC method we implemented for our analysis (see section methods for details). For each window we obtained different posteriors of the model parameters describing the RSA. In Figure 2A-B are respectively shown the log-ratio of ABC model selection score for the two models and the mixture coefficient *α* of the competition model in equation 3. Where the mixing coefficient *α* of the mixture model is higher, ABC model selection score supports the presence of competition. The results for the remaining parameters describing the sliding windows RSA are shown as Supplementary Figures B19, B20.

**Figure 2:**
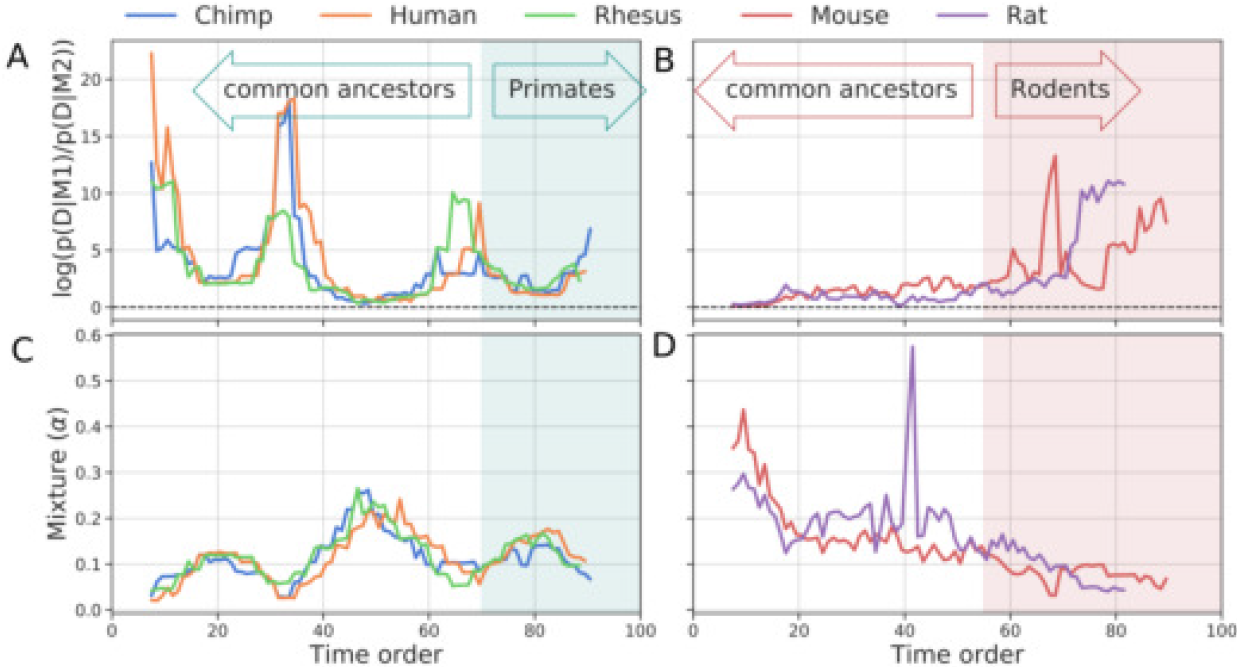
Comparison of the two model performance during the evolution of the LINEs ecosystem in primates and rodents by using the sliding window approach. Log-ratio of the acceptance ratios of the neutral and the competition models (see method section for details) are shown in panel A for primates, and B for rodents. Estimation of the mixture coefficient are shown in panel C for primates, and D for rodents. The portion of elements highlighted in different colors belong to different clusters in Figure 3.

### The variation in time of dynamics rates highlights a transition in LINEs average abundance for both rodents and primates during evolution

The study of the correlation between the negative binomial parameters *x* and ϒ obtained with the sliding window analysis of the RSA patterns is shown in Figure 3A-D. Such correlation is inherent to the parameters of the model and is related to the expectation value of the negative binomial distribution (average abundance defined in equation 5).

**Figure 3:**
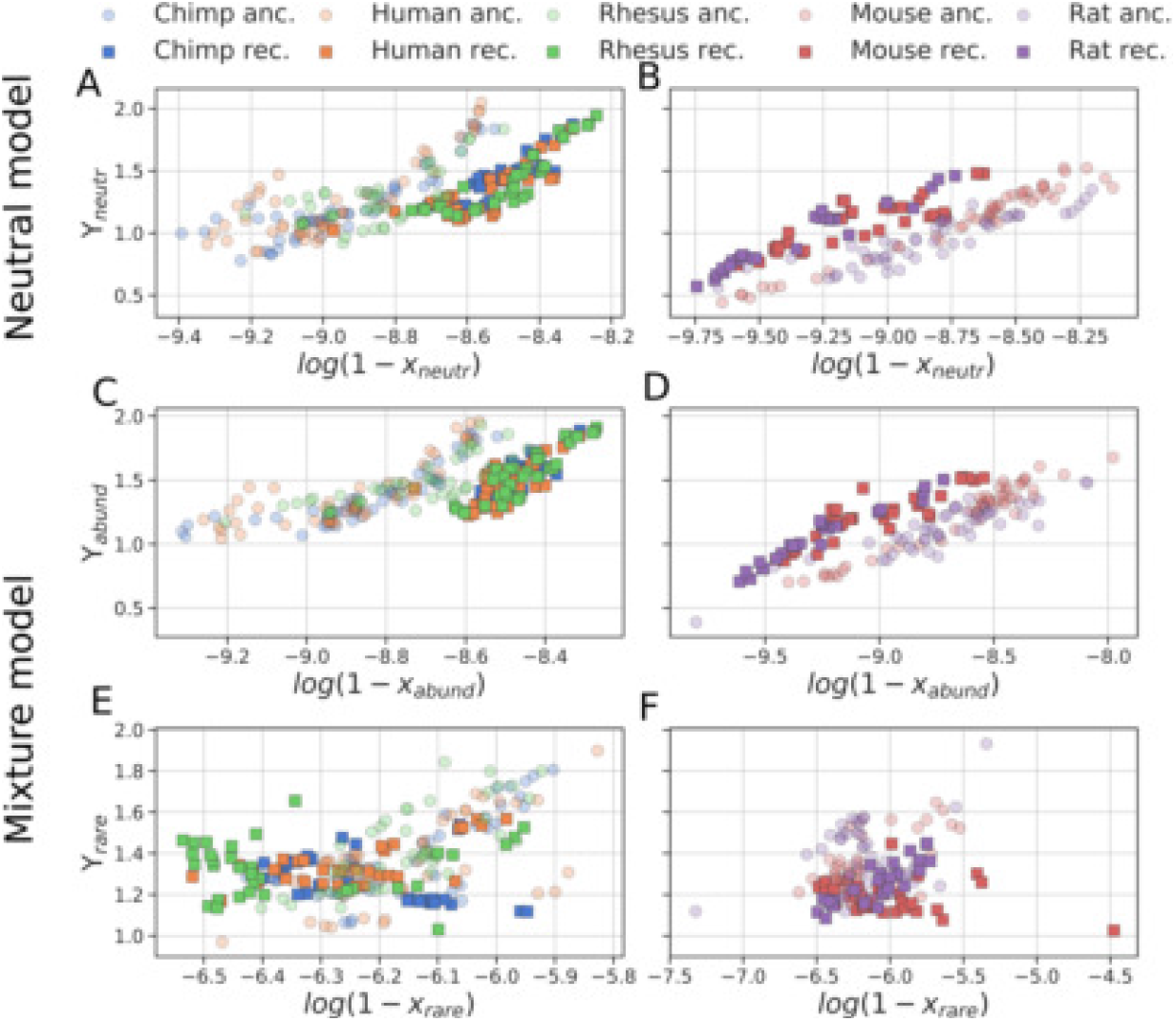
Space of parameters of the two model tested by the sliding window approach in primates and rodents. The space of parameters describing sliding window ecosystem of LINEs in human, chimpanzee, rhesus macaque (panels B,D,f)) and mouse, rat (panels A,C,e)) is shown. *x* and ϒ parameters are correlated by the expected value (mean) of the distribution. Panels A,B refer to the neutral model, panels C,D refer to the group of the mixture model with largest copy number, panels e),f) refer to the group of less abundant elements. Circles indicated the most ancient elements. Transition between the two cluster are associated with specific LINE species appearance.

The negative binomial parameters *x* and ϒ of both cohorts cluster in two groups, separated approximately at the time point associated to the appearance of specific elements (LIMA/LPB for primates and Lx for rodents). One cluster contains the results for the windows before such element appearance, the other cluster contains the windows after such element appearance. Thus, the separation between the two clusters corresponds to a transition of the average abundance of LINEs ecosystem. For the group of primates the transition is toward lower average abundance, while for the group of rodents it is toward larger average abundance.

This transition can be observed in the neutral model (Figure 3A and in the abundant copy number group (Figure 3B of the competition model, the component that includes the majority of the elements in the genome. The rare copy number group of the competition model, on the other hand, does not show this transition, and thus the two groups form a single cluster (Figure 3C. These behaviours are further commented in the discussion and Supplementary Material Section A.2.The results for the remaining parameters describing the sliding windows RSA are shown as Supplementary Figures B19, B20.)

### LINEs average abundance increase for rodents and decrease for primates; open chromatin decreases for everybody

In Figure 4 we show the sliding window analysis of the average abundance and the average percentage of insertions in open chromatin regions [45, 15]. In the primates cohort the average abundance is decreasing (Figure 4C, while in the rodents cohort it displays a fast transition to larger abundances (Figure 4D, in agreement with the results displayed in Figure 3. On the other side, the average percentage of LINE copies belonging to open chromatin regions (Figure 4A-B displays a similar decreasing temporal trend for all the organisms for which such information was available (human and mouse).

**Figure 4:**
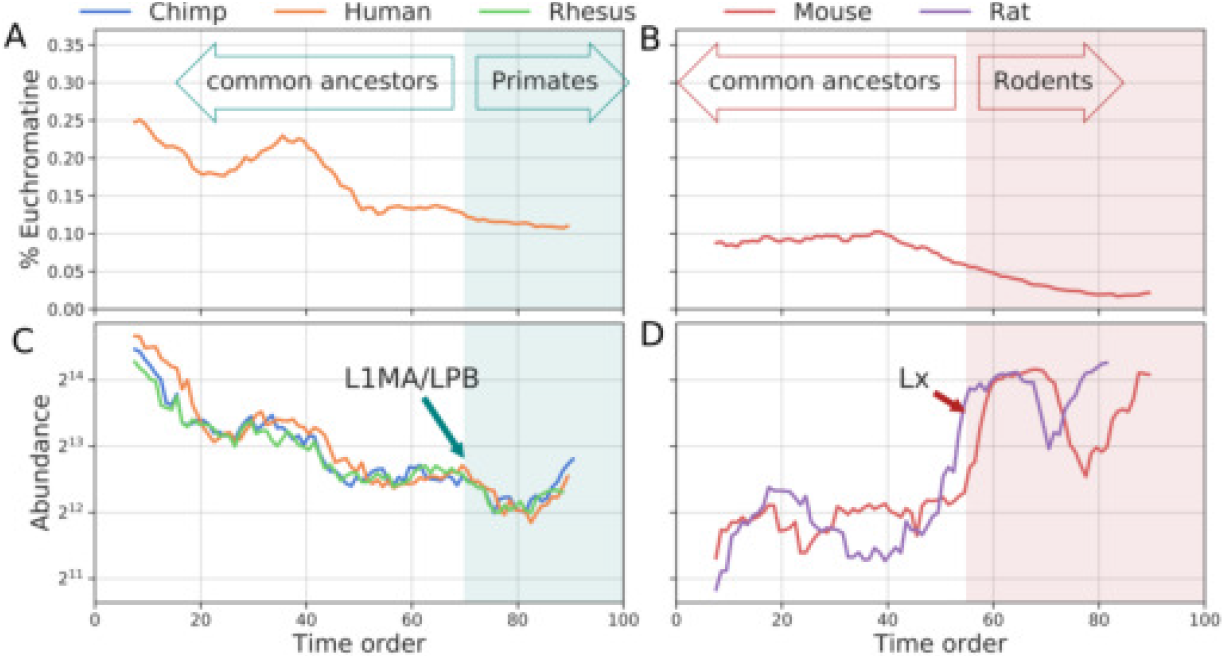
Sliding window of LINE insertions percentage inside euchromatin and expected LINEs abundance for primates and rodents cohorts. Percentage of LINE copies inserted in euchromatin regions calculated by the sliding window approach are shown in panel A for primates, and B for rodents. LINEs average copy number (sum of euchromatin and heterochromatin insertions) calculated by the sliding window approach are shown in panel C for primates, and D for rodents. The portion of elements highlighted in different colors belong to different clusters in Figure 3.

### There is a transition in LINEs chromatin configuration in rodents but not in primates

In Figure 5 we show the correlation between the number of insertions in euchromatin (ECN) and the number of insertions in heterochromatin regions (HCN), and the corresponding percentage of insertion in ECN. Two clusters are clearly visible for the mouse data set (5B,D. We applied a Gaussian mixture clustering method to separate the two clusters in mouse (see supplementary Figure B23).The elements classification in two clusters by such procedure is in strong agreement (except a few elements that fall in the wrong cluster) with a classification based on the time of appearance of the elements, with the same time threshold identified in Figure 3B,D for mouse and highlighted by color (and marker) labels in Figure 5B,D.

**Figure 5:**
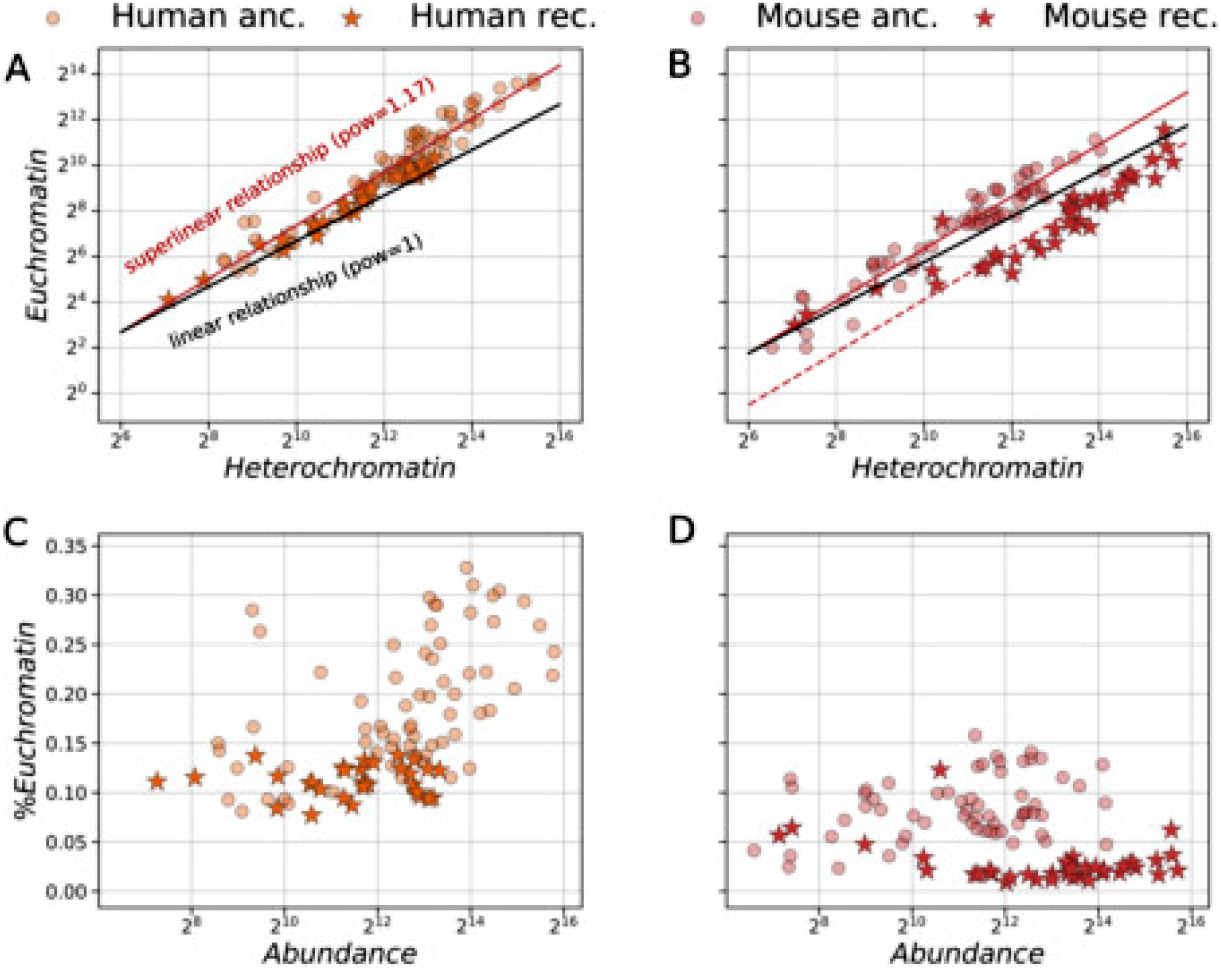
Number of insertions in euchromatin (ECN) and heterochromatin (HCN) states in human and mouse. Upper panels: Scatter plot in *log*_2_ scale of the number of insertions in euchromatin in comparison to that in heterochromatin for each LINE specie for human A and for mouse B. Trend obtained by OLS fit are shown (super linear, red lines), linear trend is also shown for comparison (black lines). Lower panels: percentage of insertions in euchromatin for human C and for mouse D. The elements highlighted in different colors belong to the two different clusters in Figure 3: ancient elements (circles), recent elements (stars).

The beginning of the transition in Figure 5B for the mouse data set is contemporary to the transition in time toward larger abundance (Figure 4D and the separation between the two clusters of parameters (Figure 3B,D for the rodents cohort. Regarding the primates cohort, in Figure 5A,C we highlighted the group of elements belonging to the cluster of the parameters containing the most recent elements (Figure 3A,C for completeness, but two clusters in terms of chromatin correlation cannot be identified (see Supplementary Figure B22).

### Larger LINE abundances are associated to a higher percentage of insertions in euchromatin regions

The logarithms of ECN and HCN shown in Figure 5 are strongly correlated. To estimate the correlation, a linear regression between the logarithm of the counts has been performed. This correlation corresponds to a power-law relationship between the raw counts:

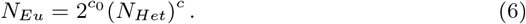

In Table 1 we display the best fit parameters, the corresponding goodness of fit and standard error of the estimated parameters obtained through Ordinary Least Squares (OLS) regression.

**Table 1:** Estimated parameters (*c* and *c*_0_) of the correlation between the number of insertions in euchromatin and the number of insertions in heterochromatin regions defined in equation 6 with their corresponding standard error (*c* SE and *c*_0_ SE) and goodness of fit (Adj. R-Squared) obtained through OLS regression of the log counts.

| Organism | $c_0 \pm \text{SD}$ | $c \pm \text{SD}$ | Adj. R-squared |
| --- | --- | --- | --- |
| Human (ancient) | $-4.3 \pm 0.4$ | $1.17 \pm 0.04$ | 0.929 |
| Human (recent) | $-3.4 \pm 0.5$ | $1.03 \pm 0.04$ | 0.968 |
| Mouse (ancient) | $-4.9 \pm 0.4$ | $1.12 \pm 0.04$ | 0.926 |
| Mouse (recent) | $-9.0 \pm 0.9$ | $1.18 \pm 0.07$ | 0.905 |

The super-linear correlation between ECN and HCN is consistent with the trend shown in Figure 5C-D, meaning that for such configuration larger LINE abundances are associated to a higher percentage of insertions in euchromatin regions for such elements.

## Discussion

The sliding window analysis of the RSA supports the presence of periods of competition between different LINEs elements during both primates and rodents evolution (Figure 2). The applied ABC method already penalizes the model with the highest number of parameters because the phase-space is larger. Hence, when the ABC model selection score approximately equal to one it supports the presence of the mixture from a statistical perspective. Furthermore, the introduction of the second component in the PDF enhance the ability to extract useful information from the data and a better characterization of the biological phenomena, for example it permits to cluster different taxonomic orders by using the corresponding rates of the dynamics (supplementary Figures B16).

The value of the failure parameter ϒ remains always closed to one (ϒ ~ 1) in both models (see Figure 3 and Supplementary Figures B19, B20),suggesting that a pure accumulation process out of equilibrium is more likely (*b_AI_* ≈ 0, *d_I_* ≈ 0). The shape of the RSA, together with the rareness of trans-complementation events, supports the idea that equilibrium between host and LINE population does not hold in general (*b_A_* ≪ *d_A_*), but a competition between the host and LINE species takes place (Supplementary Figure A2, A3). In fact, in the case of equilibrium for the active sub-type dynamics (*b_A_* ≈ *d_A_*), numerical simulations display heavier tails in the generated RSA (Supplementary Figure A3).Such excess of abundant elements could be compensated by a higher rate of excision from the genome (*d_I_*) (Supplementary Figure A3).However, this parameter configuration results less plausible, at least in primates, because more ancient LINE elements are more abundant on average than the recent one (Figure 4C).

The transition in the correlation of the parameters *x* and ϒ in Figure 3 is associated to a transition to lower average abundance for primates (Figure 3A,C) and higher average abundance for the two rodents (Figure 3B,D), as it can be noticed in equation 5). These transitions are in agreement with the trend in time of the abundance (Figure 4C,D) and further supported by the transition in the chromatin landscape of the LINE copies for mouse (Figure 5B) and by the concurrent amplification of host-specific LINE elements.

Regarding the sliding window analysis of the chromatin configuration of the copies, the percentage of LINE copies inserted in euchromatin regions displays a decreasing trend in time for both human and mouse (Figure 4A,B).

Observing the distribution of the number of copies belonging to open and closed chromatin regions for each LINE species, the super linear correlation observed in Figure 5A,B leads to the interesting result that a higher copy number, i.e., the sum of euchromatin and heterochromatin contributions, is associated to a higher percentage of insertions in euchromatin states (Figure 5C,D). A reduction of the average percentage of insertions in euchromatin regions thus corresponds to a reduction in LINEs average abundance. Moreover, the presence of the same type of correlation in human and mouse genomes, shared by all the most ancient elements, indicates the existence of a common pattern in the chromatin landscape of LINEs in mammals.

In agreement with such prediction, in primates the decreasing trend of the insertion percentage in open chromatin regions is accompanied by a decreasing trend of LINEs average abundance in time (Figure 4A,C). On the contrary, referred to mouse, the average abundance at a certain point drastically rears up (Figure 4B,D). The time point at which the abundance rears up corresponds, in Figure 5B, to a transition to a smaller value of the coefficient *c*_0_ (see Table 1), while for ancient LINE species the correlation trend in mouse genome is very close to the one observed in human. Given the same number of insertions in open chromatin regions, a lower value for *c*_0_ corresponds to a larger abundance, and, consequently, to a lower percentage of insertions in euchromatin.

This relationship between average abundance and chromatin state can be interpreted from the perspective of the host organism fitness. A transition to a higher average copy number (Figure 4D) may have a negative impact on the host fitness if it is not compensated by some regulation mechanisms, because this would increases the probability of deleterious insertions in the genome. The combined transitions in the chromatin landscape and abundances, further associated to the evolutive transition of the host genome (murine subfamily differentiation), observed in mouse, support this idea, and are perhaps the result of the competition between the host and LINEs [9].

In fact, the transition in mouse to a different chromatin configuration of the LINEs insertions (depicted in Figure 5B,D) is related to the time of appearance of the oldest element belonging to the cluster of recent elements (color labeled in Figure 5B,D). Such transition is contemporary to the transition to higher average LINEs abundance as shown in Figure 3B,D and Figure 4D, and corresponds to the appearance of the LINE family Lx. The amplification of the LINE family Lx is associated with the murine subfamily evolutive radiation that happened ~ 12 Myr ago [30, 16]. In fact, the elements belonging to the cluster of the recent elements are mainly murine specific.

Instead, the chromatin configuration of the LINEs insertions in human (depicted in Figure 5A,C does not present a transition between two different chromatin state distributions, contrary to what observed for mouse. In human a transition to a lower average copy number is observed (Figure 3A,C and 4C). This could be the reason why chromatin states distribution in human is not affected significantly, since further changes were not necessary to preserve the host fitness. Indeed, the most ancient LINE species involved in the transition depicted in Figure 3A,C are related to the evolutive differentiation of Primates, associated with the amplification of LIMA/LPB subfamilies ~ 70 − 100 Myr ago [26].

## Conclusion

The present research is a first attempt to discover evolutionary footprints of transposons activity on genomes by merging bioinformatics and biophysics results obtained from different data types, with the final objective to give indication for discovering and/or confirmation in wet labs.

The variation in time of the chromatin state of LINE copies together with the average abundance and the estimated rates of activity should reflect the interaction of LINE species with the host, by mechanisms of silencing (for example methylation) and self-regulation (for example selection of elements with lower birth rates or with specific genomic region preference of the new insertions). In fact, the variation in time of the estimated rates of activity, combined with the average abundances and chromatin state of LINEs copies, displays evidences of host-elements interaction and features highlighting taxa-specific element appearance, such as Lx, associated to the radiation of the murine subfamily, and LIMA/LPB sub-families, associated to primates evolution. This is in agreement with several independent studies [26, 38, 29, 30]. Our results therefore suggest the existence of a general mechanism of chromatin response to the increase of transposon activity.

This hypothesis could be further investigated by considering the chromatin structure of other organisms to verify consistency with the obtained results (in the present cohort Rhesus and Chimp for primates and Rat for rodents). Such experiments could benefit also from the integration of molecular information at the copy level, such as DNA sequence, chromosome location and genome patterns (CG/AT content, chromatin configuration, etc…), which is available for many organisms nowadays.

We surmise that the interdisciplinary approach of combining general biophysics models for genomics elements with molecular information at copy level could become a powerful tool to understand present and past general dynamics of the entities that contribute in shaping the genome, and lead to new insights bridging evolutionary biology, genome ecology, and transposon ecology fields.

## Acknowledgements

The results presented in this work are part of the PhD Thesis of SV [43]. This work has been supported by the Italian Ministry of Education at University of Bologna (Alma Mater Studiorum), Department of Physics and Astronomy (DIFA), by the Basque Center for Applied Mathematics (Bilbao, Spain) and in part by the following NIH grants: R56 AG050582-01 to N.N. and F31AG050365 to S.W.C‥ S.W.C. was also supported by the NIH Institutional Research Training Grant T32 GM007601. We also aknowledge IMforFuture EU project and HARMONY EU project.

## Author contributions statement

N.N. and G.C. conceived the study, S.W.C. and I.F.V. prepared the data sets, E.G., S.V. and C.S. implemented the ABC pipeline and dynamics simulations, S.V. performed the analysis. All the authors contributed to the development of the study. S.V., E.G., C.S. and G.C. wrote and revised the manuscript.

